# Reliable one-step assessment of IGHV mutational status and gene mutations in Chronic Lymphocytic Leukemia by capture-based high throughput sequencing

**DOI:** 10.1101/2022.03.09.483581

**Authors:** Yannick Le Bris, Florian Thonier, Audrey Menard, Olivier Theisen, Béatrice Mahe, Anne Lok, Simon Bouzy, Marie C Béné

**Affiliations:** Hematology Biology, CHU de Nantes, Nantes, France; Inserm 1232, CRCINA Université de Nantes, Nantes, France; INRIA, Rennes, France; Hematology Clinic, CHU de Nantes, Nantes, France

**Author notes:** **Contact information** Dr Yannick Le Bris. **Competing interests.** None.

## Abstract

Proper management of chronic lymphocytic leukemia (CLL) patients requiring therapy relies on two important prognostic and theranostic molecular features: respectively, the mutational status of tumoral cells immunoglobulin heavy chain variable domain (*IGHV)* and the characteristics of *TP53*. Both these (immuno)genetic analyses require multiple time-consuming amplification and sequencing techniques by Sanger or HTS. The capture-HTS technology, allowing to select regions of interest, represents an attractive alternative and has already been applied for the detection of clonality in lymphoproliferative disorders. Here, a single-step capture design was developed to concomitantly investigate for *IGHV* and *TP53.* This was applied to a training retrospective (n=14) and a validation prospective (n=91) cohorts of CLL patients. The training cohort demonstrated the robustness of the method by comparison with the classical Sanger sequencing technology (100% identical results) for the *IGHV* mutational status. This consistency was confirmed for the first 59 patients of the validation cohort. Overall, the *IGHV* status of whole population (n=103) was accurately identified. Simultaneously, deletion or mutations of *TP53* were identified from the same capture-library and HTS-sequencing run for each patient. This novel approach provides, in a single assay, useful answers about the molecular landscape of CLL patients, allowing for a documented choice of therapy.

**Graphical abstract:** 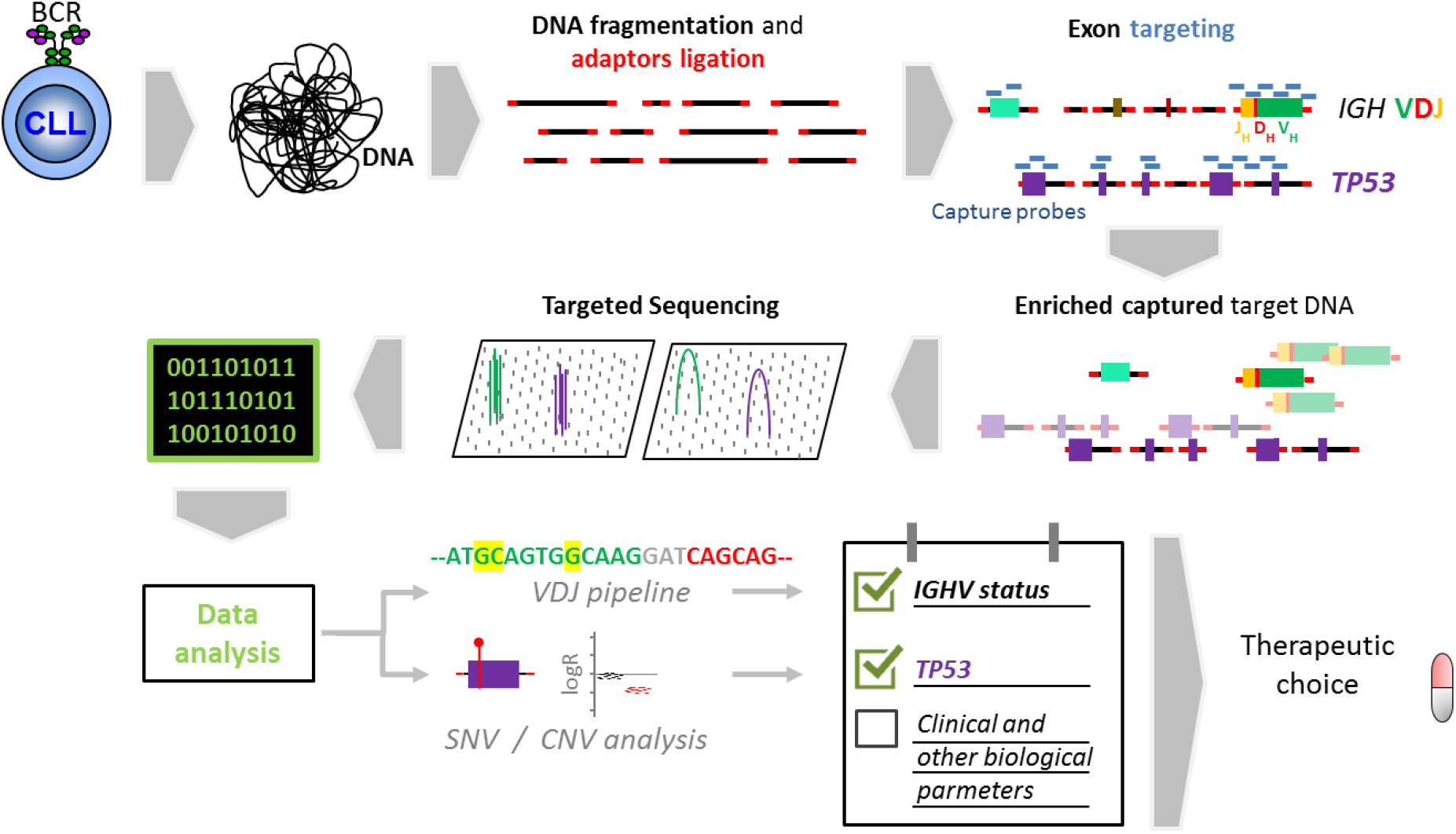

## Introduction

An impressive array of new strategies has recently been developed for patients with chronic lymphocytic leukemia (CLL) having reached a stage requiring therapeutic management (1). Two major molecular characteristics of the disease now guide the best treatment schedule’s choice. One investigates the tumor suppressor protein TP53, the alleles of which being eventually deleted or mutated (2). This feature is important to check both before first line treatment and at relapse as it is susceptible to be modified by treatment pressure favoring the emergence of subclones (3). The other one, stable over time, is the rearrangement status of the variable domain of the immunoglobulin heavy chain gene (*IGHV*) of the clonal tumor cells (4). Indeed, an unmutated status has been demonstrated to be an important poor prognosis marker and may now lead to a better therapeutic choice in first line treatment. Other high-risk markers such as complex karyotype (5,6), Bruton tyrosin kinase (BTK) C81S mutation or phospholipase C-gamma (*PLCG2*) variants are also useful to guide therapeutic choices (7,8). The management of CLL patients therefore now relies even more on a close interaction between clinical and laboratory hematologists, the latter being expected to provide the necessary information for appropriate treatment choice.

The European Research Initiative on Chronic Lymphocytic Leukemia (ERIC) (www.ericll.org) has provided recommendations on methodological approaches for *TP53* mutation and *IGHV* mutational status analysis (9). Technically, the use of high throughput sequencing (HTS) is favored for the identification of mutated *TP53* subclones (10). HTS has also recently been proposed for the estimation of *IGHV* mutational status with an amplicon strategy (11). In practice, fulfilling current laboratory requirements in order to adapt therapeutic choices in CLL has become a painstaking, time consuming succession of multiple PCRs with various primer sets, followed by Sanger or HTS sequencing and bioinformatics analyses.

We propose here an innovative one-step HTS-capture sequencing approach that allows for an exhaustive evaluation of *IGHV* mutational status meeting the technology requirements recommended by the ERIC Group. Moreover, this is combined in a single robust step with investigation of the presence or absence of *TP53* and other oncogenes variants of theranostic interest. This approach is shown here in two cohorts of training and validation with successful comparison with classical time-consuming techniques.

## Materials and methods

### Patients

Overall, samples from 103 patients were used in this study. In order to validate the capture assay for *IGHV* mutational satus, 14 patients (training cohort) were selected retrospectively as having benefited from a Sanger analysis as pre-treatment assay. A confirmatory cohort of 91 patients, addressed for pre-therapeutic *IGHV* assessment, was then tested prospectively. DNA (n=103) and/or RNA (n=28) were extracted from peripheral blood (PBMC) (n=97) or bone marow mononuclear cells (n=4) or from sorted CD19+ peripheral cells (n=2). All patients provided informed consent and the study was conducted according to good medical and laboratory practice and French legislation.

### IGHV assessment using Sanger historical method

For comparison with capture HTS results, the gold standard for the *IGHV* status analysis was Sanger sequencing with VH Leader (n=24) and/or IGH Biomed-2 PCR primers (n=31) (12) according to ERIC recommendations (13,14). In the training cohort, clonal *IGHV* rearrangements were detected by fragment analysis then Sanger sequenced. In the confirmatory cohort, Sanger sequencing was performed on a selection of patients with primers chosen on the basis of the HTS reads. IMGT (15) and Arrest (16) databases were used in all cases for sequence interpretation according to ERIC recommendations (17).

### Probes design and high throughput sequencing (HTS)

The Agilent Sureselect XTHS2 technology was used. Capture probes, of a 120 bp size, were designed to target functional sequences of variable (V), diversity (D) and junctional (J) segments of the *IGH* gene and of *TP53, BTK, PLCG2, BRAF, NRAS, KRAS* and *MYD88*. These probes were designed based on genomic hG38 positions proposed by the IMGT database (18). Some *IG* segments absent from the hG38 haplotype were also included (19). Capture was performed with 50ng of DNA per patient after a fragmentation time of 5 minutes, followed by 10 PCR cycles. A MiSeq (Illumina) instrument was used for sequencing with 500V (n=69) or 600V2 (n=34) cartridges.

### Bioinformatics

#### Initial VDJ detection and clustering with the Vidjil-algo

VDJ recombinations were identified through <www.vidjil.org>. This algorithm selects reads with both enough V *k-mers* (subsequences), then enough J k-mers, identifying a *window* with a default length of 60bp centered on the relevant complementary determining region 3 (CDR3). In the next step, it clusters the reads into *clonotypes* sharing the exact same window, thus building a consensus sequence (20). Some reads are filtered out because they do not show enough V and J k-mers. The latter include non-recombined sequences, but also actual V(D)J recombinations that are *shifted* by a few nucleotides in the 5' or 3' direction, but caught by the capture protocol where sequencing starts at a random position.

#### Read identification and consensus sequence extension

To extend clonotypes consensus sequences for a better identification of V genes and evaluate hypermutations, post-processing was performed by re-using some of the filtered-out reads. The selection of reads from other recombinations or non recombined reads was avoided by retaining, for each clonotype, reads sharing again the exact 60bp window sequence on either the sense or antisense sequence. The *de novo* assembler (21)*Minia* was used to consolidate the extended clonotypic reads (both initial and reused) and exclude sequences from minor clonotypes. Built contigs and consensus sequences were examined without any reference sequences. Only consensus sequences longer that the ones identified after initial analysis with the Vidjil-algo were considered and used to compute the hypermutation status.

Of note, this post-process is available in open-source on <https://gitlab.inria.fr/vidjil/contrib> and can be activated on any Vidjil server.

#### Variants and CNV

The SeqOne^®^ software (22) was used for SNV and CNV analysis of the oncogenes sequenced. Quality parameters considered were a minimum of 200X coverage on coding sequences of interest, Q30>96% and 1% or more of variant allele frequency (VAF).

## Results

### Comparison of Sanger and capture

The training cohort was used to evaluate the comparability between the gold standard Sanger sequencing and the novel capture method. For the 14 selected samples, as mentioned above, the V family of interest had been identified by a clonality analysis based on the Biomed primers and methodology (12). The V family detected was then used for amplification and Sanger sequencing. Finally, sequence comparison was performed with the IMGT algorithm (15) to determine the mutational status of each clone.

The one-step capture assay successfully identified simultaneously the size of the clone, its V family and its percentage of homology with the germline sequence, i.e. mutational status by direcly consulting the IMGT database within the Vijil application.

The comparison was of 100% concordance in determination of the *IGHV* mutational status in these 14 cases.

For further confirmation of the capture method’s robustness, the V family identified by capture was used for amplification and Sanger sequencing in the first 39 samples from the validation cohort. As shown in Figure 1, after sequencing and IMGT analysis, linear regression yielded a strong correlation (r^2^=0.99; p<0.001) of mutational status percentages between both methods. Of note, HTS revealed one case that had two productive clonotypes with different mutational statuses (%homology: 95,14% and 100% respectively). Only the mutated clonotype (%homology: 95,14%) had been detected by PCR clonality analysis. Three patients could not be analyzed by Sanger sequencing due to sequence mixing (n=2) or sequencing failure with VHL and Biomed-2 primers (n=1). Overall, the *IGHV* sequence could not be determined with the traditional PCR approach for 4 cases out of 53 tested. For these 4 cases the sequence and therefore mutational status of the productive IGH clone was successfully determined by HTS.

**Figure 1.**
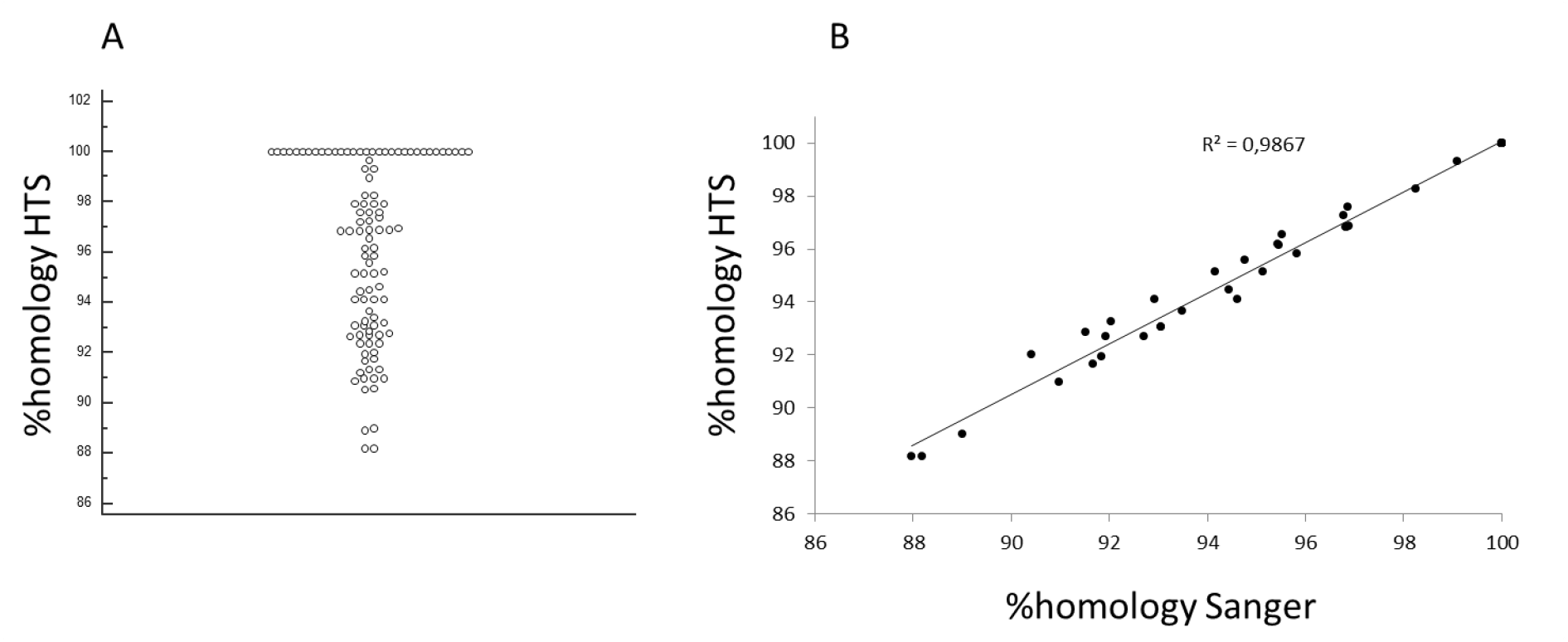
Distribution (A) and comparative (B) analysis of the homology percentages of IGHV segments sequenced by HTS *versus* Sanger.

### Capture-based IGHV analysis

Considering both cohorts, 103 patients (including 97 CLL and 6 small lymphocytic lymphoma with a circulating phase) addressed for pre-therapeutic *IGHV* mutational status assessment were analysed. They were 69 males and 34 females, with a median age of 71 years (range: 34-89).

HTS clonality analysis allowed to obtain a median of 649 reads (range 132-1770) with detectable *IGH* rearrangements. The median consensus reads size of these productive IGH clonotypes was 255 bp (120-334) before contiguity assessment. Realignement subsequently allowed to increase reads lengths up to a median of 709 bp (381-989), allowing an analysis of much more than the required 250 bp of the IGHV complete sequence in all cases.

The median depth of analyzed productive tumoral clonotypes was 247 reads (54-1759). Four different IGH clonal profiles were disclosed, respectively i)one complete VDJ plus one partial DJ (n=79), ii)one productive VDJ plus one unproductive VDJ (n=13), iii)only one productive VDJ (n=9) and iv)	two productive VDJ plus one or two partial DJ (n=2).

Considering productive tumoral rearrangements, as expected, a selection bias was observed for IGHV families with a majority of VH3 and 48% of IGHV segments represented by only 6 different segments (IGHV1-69, IGHV4-34, IGHV1-2, IGHV3-21, IGHV3-23, IGHV3-48) (Figure 2A, B). There were 12% of stereotyped CDR3s among productive IGH rearrangements (Figure 2C).

**Figure 2.**
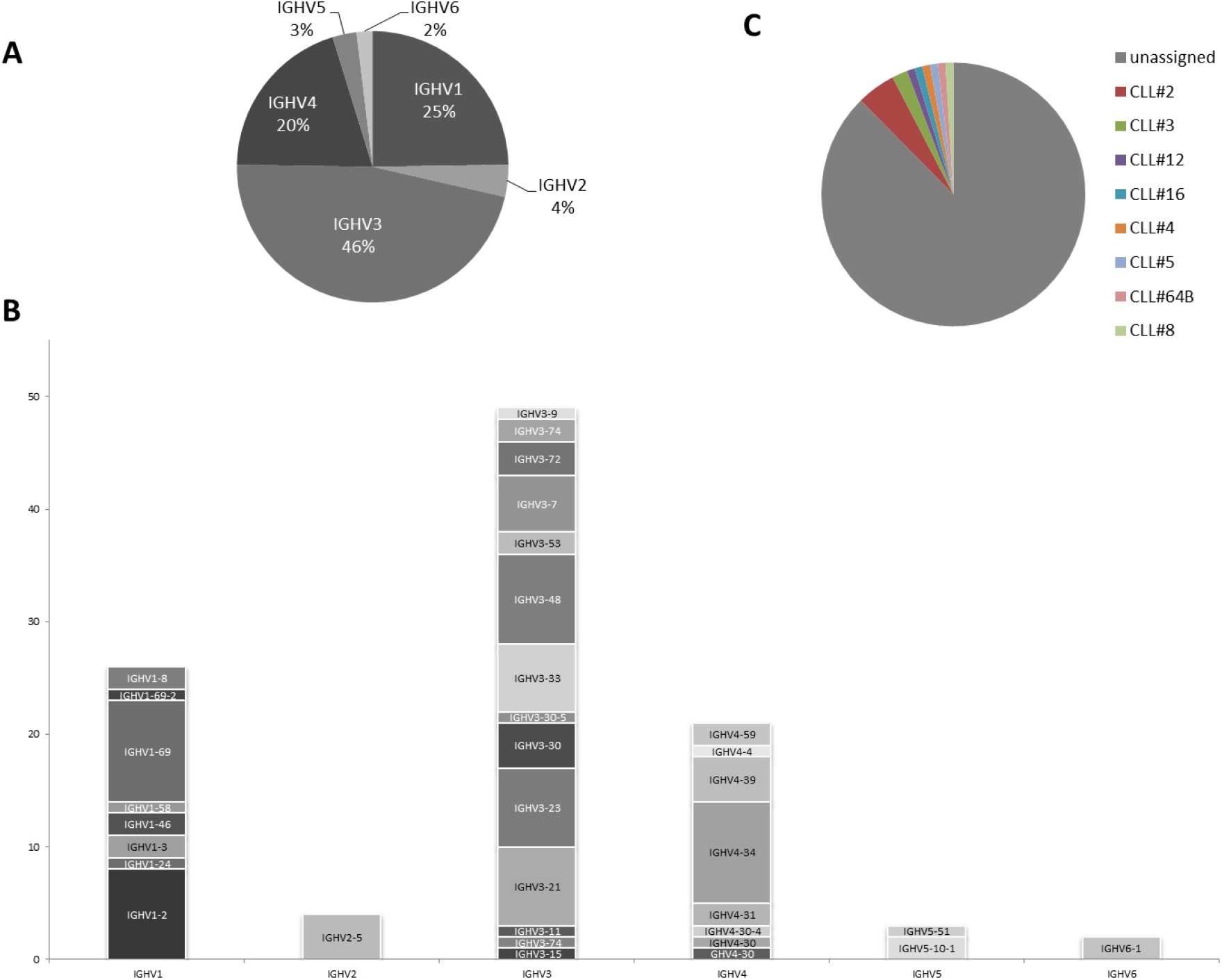
Profile of IGHV segments identified in HTS. a. Distribution of HV families detected by NGS. B. Distribution of *IGHV* segments identified for each of the VH families. C. Distribution of the identified subsets among the detected IGH segments.

Clonality and the mutational status were demonstrated and identified in all cases, fulfilling ERIC requirements. Global distribution of the homology percentage of the productive IGH rearrangements detected with the HTS approach showed 38 cases with unmutated IGHV (homology>98%) and 67 mutated cases (homology<98%). Of note, as shown avbove, two patients presented with two productive rearrangements.

### Oncogene variants analysis

Analysis of the sequenced capture genes identified mutations in 35 patients (Figure 3). Thirty nine *TP53* mutations were detected in 16 patients (16%) with a median of 1 mutation per patient (1-15) and a median allele frequency of 4% (1-90). The patient who had 15 *TP53* mutations did not display CNA for this gene. Mutations were also observed in *NRAS* (8%), *BRAF* (4%), *MYD88* (3%) and *KRAS* (2%), but none was found for *BTK* or *PLCG1*. *KRAS* gains were found in 8% of the cases. Finally, duplication of an unmutated *BTK* gene was disclosed in one patient with lymphocytic lymphoma for whom a first-line pre-therapeutic workup was performed due to nodal progression.

**Figure 3.**
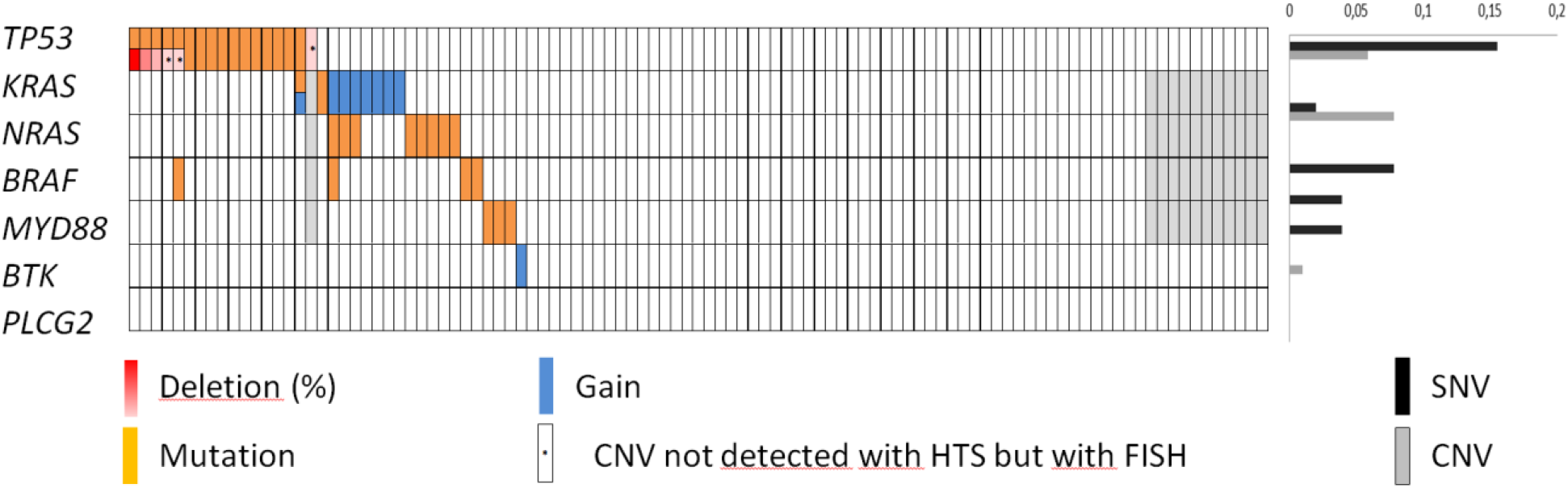
Profiles of mutation (SNV) and of copy number variation (CNV) detected among the oncogenes studied.

The performance for CNVs detection was similar between HTS capture and fluorescence in situ hybridization (FISH) 68 of 70 cases tested. These were *TP53* deletions in 4 cases. The three discordant cases were *TP53* heterozygous deletions in respectively 10, 10 and 11 percent of the nuclei in FISH, not detected in HTS (Figure 3). However, two of these cases displayed *TP53* mutations identified in HTS.

## Discussion

This report presents the performance of an innovative single-step combined analysis of *IGHV* mutational status and oncogene somatic variants of therapeutic interest, in the context of CLL, by capture HTS. Identification of the major clone was effective in all cases, including the V, D and J families involved, full CDR3 sequence and mutational status. Moreover, the same run explored oncogenic mutations of potential impact. This comprehensive and robust strategy could considerably lighten the workload and reduce the turnaround time currently necessary to provide this information for proper CLL patients management.

An amplicon-based PCR strategy remains the gold-standard to identify *IGHV* rearrangement of CLL clones. Sanger sequencing has been applied historically and, more recently, HTS methods have been developed, still following PCR amplification (11). In some cases, however, a proper identification of the malignant clone is hampered by somatic hypermutations (SHM) impairing the hybridation of amplification primers. This is especially the case with some of the leader *VH* primers recommended by ERIC (13,14). Another hindrance is observed with some sequencing instruments, unable to produce long enough reads not covering the complete *VDJ* sequence (23). In the past few months, the use of innovative tools for immunoglobulin genes analyses has been reported, such as the nanopore strategy, demonstrated as a good alternative for *IGHV* status assessment in CLL (24). Capture HTS has also been shown recenlty by the EuroClonalty group as an efficient means to assess clonalty in B-lineage lymphoproliferative disorders (25).

Capture enrichment relies on a different strategy, based on a first step of fragmentation of the patient's genomic DNA. DNA fragments are then selected by hybridizing to 120 bp complementary sequences of interest. DNA fragments that do not hybridize are thus removed. A short PCR amplification, of ten cycles, allows to increase the quantity of the sequences retained and degrade the capture probes. Sequencing is then carried out on a high-throughput sequencer. The difference with the amplicon approach is that amplification defects (when the PCR primer is on a mutation) are avoided because the capture probe will bind the region anayway and moreover will not be hampered by SHM. In addition, the lower number of PCR cycles in capture reduces sequencing errors.

Because capture can be designed to encompass many genome sequences, we figured that in addition to the detection of clonality, it could also be applied to a full comprehensive sequencing also allowing for the assessment of mutational status. The originality of this report lies in the combined strategy of a large number of capture probes and post-sequencing contiguity alignment, yielding to the obtention of very long sequences, largely over the ERIC’s required 250bp and specifically covering the entire *V-D-J* region of the rearranged *IGH* gene. This extensive coverage allows for a reliable sequence analysis and limits the risk of identifying pseudo-mutations related to sequencing artifacts. Indeed, similar mutational results were obtained for 100% of the patients evaluable with both capture-HTS and PCR/Sanger, validating the reliability of this new technological approach. IGHV mutational status analysis was furthermore successful in all 105 patients studied with HTS-capture while the traditional PCR approach failed in 4/59 cases (7%) tested with both methods. Surprisingly, a much higher frequency (79%) of partial DJ rearrangements of the IGH locus was observed than in previously published data with an amplicon technique (32%) (26). This difference could be related to the presence of SHMs that made difficult to amplify partial DJ rearrangements in former studies. Indeed, capture probes ignore SHMs and allow proper sequencing of even partial rearrangements. The potential hindrance is that this methodology is likely to capture pseudogenes with high homology with functional genes, leading to off-target sequencing (about 30%, data ot shown).

This report also demonstrates the simultaneous detection, with the capture approach, of potential anomalies of *TP53* and other genes with high sensitivity in the same run. This is an important advantage in patient management by providing in one-step all necessary information to decide for the optimal therapeutic strategy. Indeed, *IGHV* and *TP53* mutational status are currently the major theranostic markers in CLL, orienting treatment towards immunochemotherapy or chemo-free (BTKi, iBCL2) schedules (2,27,28). It would moreover be very easy to include other targets such as *ATM, NOTCH1, SF3B1, BIRC3* (29) or others, should they become significant indicators of response or prognosis.

In summary, the molecular analysis strategy depicted in this study and the consistently robust results obtained provide an interesting new way to swiftly explore pronostic and theranostic markers for CLL patients requiring treatment. On a broader perspective, the possibility of analyzing the complex *IG* repertoire with full coverage, without the inconvenients of whole exon sequencing, is an enticing approach that could be applied both to the exploration of other B-lineage LPDs.

## Acknowledgements

The authors would like to thank Laetitia Aubert, Marie-Christine Boursier, Emilie Brangoulo, Cécile Lagarde, Veronique Chenais and Amandine Sebie for excellent technical expertise on molecular analyses. We also thank François Lozach (Agilent) for his help in designing the capture probes.

## References

1. Gaidano G, Rossi D. The mutational landscape of chronic lymphocytic leukemia and its impact on prognosis and treatment. Hematology Am Soc Hematol Educ Program. 2017;2017:329–337. Hallek M. Chronic lymphocytic leukemia: 2020 update on diagnosis, risk stratification and treatment. Am J Hematol. 2019;94(11):1266–1287. doi: 10.1002/ajh.25595. Epub 2019 Oct 4. PMID: 31364186.

2. Hallek M, Cheson BD, Catovsky D, Caligaris-Cappio F, Dighiero G, Döhner H, et al. iwCLL guidelines for diagnosis, indications for treatment, response assessment, and supportive management of CLL. Blood. 2018;131:2745–2760.

3. Rossi D, Khiabanian H, Spina V, Ciardullo C, Bruscaggin A, Famà R, et al. Clinical impact of small TP53 mutated subclones in chronic lymphocytic leukemia. Blood. 2014; 123:2139–2147. doi: 10.1182/blood-2013-11-539726. Epub 2014 Feb 5. PMID: 24501221; PMCID: PMC4017291.

4. Parikh SA, Strati P, Tsang M, West CP, Shanafelt TD. Should IGHV status and FISH testing be performed in all CLL patients at diagnosis? A systematic review and meta-analysis. Blood. 2016; 127:1752–1760.

5. Le Bris Y, Struski S, Guièze R, Rouvellat C, Prade N, Troussard X, et al. Major prognostic value of complex karyotype in addition to TP53 and IGHV mutational status in first-line chronic lymphocytic leukemia. Hematol Oncol. 2017; 35:664–670.

6. Kittai AS, Miller C, Goldstein D, Huang Y, Abruzzo LV, Beckwith K, et al. The impact of increasing karyotypic complexity and evolution on survival in patients with CLL treated with ibrutinib. Blood. 2021; 138:2372–2382.

7. Woyach JA, Furman RR, Liu TM, Ozer HG, Zapatka M, Ruppert AS, et al. Resistance mechanisms for the Bruton's tyrosine kinase inhibitor ibrutinib. N Engl J Med. 2014; 370:2286–2294.

8. Quinquenel A, Fornecker LM, Letestu R, Ysebaert L, Fleury C, Lazarian G, et al. French Innovative Leukemia Organization (FILO) CLL Group. Prevalence of BTK and PLCG2 mutations in a real-life CLL cohort still on ibrutinib after 3 years: a FILO group study. Blood. 2019;134:641–644.

9. Malcikova J, Tausch E, Rossi D, Sutton LA, Soussi T, Zenz T, et al.; European Research Initiative on Chronic Lymphocytic Leukemia (ERIC) — TP53 network. ERIC recommendations for TP53 mutation analysis in chronic lymphocytic leukemia-update on methodological approaches and results interpretation. Leukemia. 2018; 32:1070–1080.

10. Bomben R, Rossi FM, Vit F, Bittolo T, D’Agaro T, Zucchetto A et al. TP53 Mutations with low variant allele frequency predict short survival in chronic lymphocytic leukemia. Clin Cancer Res. 2021; 27):5566–5575.

11. Davi F, Langerak AW, de Septenville AL, Kolijn PM, Hengeveld PJ, Chatzidimitriou A, et al.; ERIC, the European Research Initiative on CLL, and the EuroClonality-NGS Working Group. Immunoglobulin gene analysis in chronic lymphocytic leukemia in the era of next generation sequencing. Leukemia. 2020; 34:2545–2551.

12. van Dongen JJ, Langerak AW, Brüggemann M, Evans PA, Hummel M, Lavender FL, et al. Design and standardization of PCR primers and protocols for detection of clonal immunoglobulin and T-cell receptor gene recombinations in suspect lymphoproliferations: report of the BIOMED-2 Concerted Action BMH4-CT98-3936. Leukemia. 2003; 17:2257–2317.

13. Ghia P, Stamatopoulos K, Belessi C, Moreno C, Stilgenbauer S, Stevenson F, et al.; European Research Initiative on CLL. ERIC recommendations on IGHV gene mutational status analysis in chronic lymphocytic leukemia. Leukemia. 2007; 21:1–3.

14. Huet S, Bouvard A, Ferrant E, Mosnier I, Chabane K, Salles G, et al. Impact of using leader primers for IGHV mutational status assessment in chronic lymphocytic leukemia. Leukemia. 2020; 34:2257–2259.

15. Brochet X, Lefranc MP, Giudicelli V. IMGT/V-QUEST: the highly customized and integrated system for IG and TR standardized V-J and V-D-J sequence analysis. Nucleic Acids Res. 2008; 36 (Web Server issue):W503–8.

16. Bystry V, Agathangelidis A, Bikos V, Sutton LA, Baliakas P, Hadzidimitriou A, et al.; European Research Initiative on CLL. ARResT/AssignSubsets: a novel application for robust subclassification of chronic lymphocytic leukemia based on B cell receptor IG stereotypy. Bioinformatics. 2015; 31:3844–3846.

17. Rosenquist R, Ghia P, Hadzidimitriou A, Sutton LA, Agathangelidis A, Baliakas P, et al. Immunoglobulin gene sequence analysis in chronic lymphocytic leukemia: updated ERIC recommendations. Leukemia. 2017; 31:1477–1481.

18. Giudicelli V, Chaume D, Lefranc MP. IMGT/GENE-DB: a comprehensive database for human and mouse immunoglobulin and T cell receptor genes. Nucleic Acids Res. 2005; 1,33(Database issue):D256–61.

19. Watson CT, Steinberg KM, Huddleston J, Warren RL, Malig M, Schein J, et al. Complete haplotype sequence of the human immunoglobulin heavy-chain variable, diversity, and joining genes and characterization of allelic and copy-number variation. Am J Hum Genet. 2013; 92:530–546.

20. Giraud M, Salson M, Duez M, Villenet C, Quief S, Caillault A, et al. Fast multiclonal clusterization of V(D)J recombinations from high-throughput sequencing. BMC Genomics. 2014; 15:409

21. https://github.com/GATB/miniaSeqone Last accesses Fabruary 22, 2022

22. Boulland ML, Vic S, Thonier F, Ganard M, Lamy T, Fest T, et al. Reliable IGHV status assessment by next generation sequencing in routine practice for chronic lymphocytic leukemia. Leuk Lymphoma. 2021; 1:1–4.

23. Minervini CF, Cumbo C, Redavid I, Conserva MR, Orsini P, Zagaria A, et al. Nanopore sequencing approach for immunoglobulin gene analysis in chronic lymphocytic leukemia. Sci Rep. 2021; 11:17668.

24. Stewart JP, Gazdova J, Darzentas N, Wren D, Proszek P, Fazio G, et al.; EuroClonality-NGS Working Group. Validation of the EuroClonality-NGS DNA capture panel as an integrated genomic tool for lymphoproliferative disorders. Blood Adv. 2021 ;5:3188–3198.

25. Tsakou E, Agathangelidis A, Agathagelidis A, Boudjoghra M, Raff T, Dagklis A, et al. Partial versus productive immunoglobulin heavy locus rearrangements in chronic lymphocytic leukemia: implications for B-cell receptor stereotypy. Mol Med. 2012; 18:138–145.

26. Quinquenel A, Aurran-Schleinitz T, Clavert A, Cymbalista F, Dartigeas C, Davi F, et al. Diagnosis and treatment of Chronic Lymphocytic Leukemia: Recommendations of the French CLL study group (FILO). Hemasphere. 2020; 4:e473.

27. Hallek M, Al-Sawaf O. Chronic lymphocytic leukemia: 2022 update on diagnostic and therapeutic procedures. Am J Hematol. 2021; 96:1679–1705.

28. Albiol N, Arguello-Tomas M, Moreno C. The road to chemotherapy-free treatment in chronic lymphocytic leukaemia. Curr Opin Oncol. 2021; 33(6):670–680..

29. van der Straten L, Hengeveld PJ, Kater AP, Langerak AW, Levin MD. treatment approaches to chronic lymphocytic leukemia with high-risk molecular features. Front Oncol. 2021; 11:780085.

